# Circadian Control of Heparan Sulfate Levels Times Phagocytosis of Amyloid Beta Aggregates

**DOI:** 10.1101/2021.05.04.442651

**Authors:** Gretchen T. Clark, Yanlei Yu, Cooper A. Urban, Guo Fu, Chunyu Wang, Fuming Zhang, Robert J. Linhardt, Jennifer M. Hurley

## Abstract

Alzheimer’s Disease (AD) is a neuroinflammatory disease characterized partly by the inability to clear, and subsequent build-up, of amyloid-beta (Aβ). Aβ clearance is regulated by several pathways and has a circadian component. However, the mechanism underlying the circadian clearance of Aβ has not been defined. Myeloid-based phagocytosis, a key mechanism in the metabolism of Aβ, is circadianly-regulated, presenting a potential mechanism for the circadian clearance of Aβ. In this work, we revealed that the phagocytosis of Aβ42 undergoes a daily oscillation that is dependent on the circadian clock. We found the circadian timing of global heparan sulfate proteoglycan (HSPG) biosynthesis was the molecular timer for the clock-controlled phagocytosis of Aβ and that both HSPG binding and Aβ42 aggregation were essential for this oscillation. These data highlight that circadian regulation in immune cells may play a role in the intricate relationship between the circadian clock and AD.

**Graphical Abstract:** 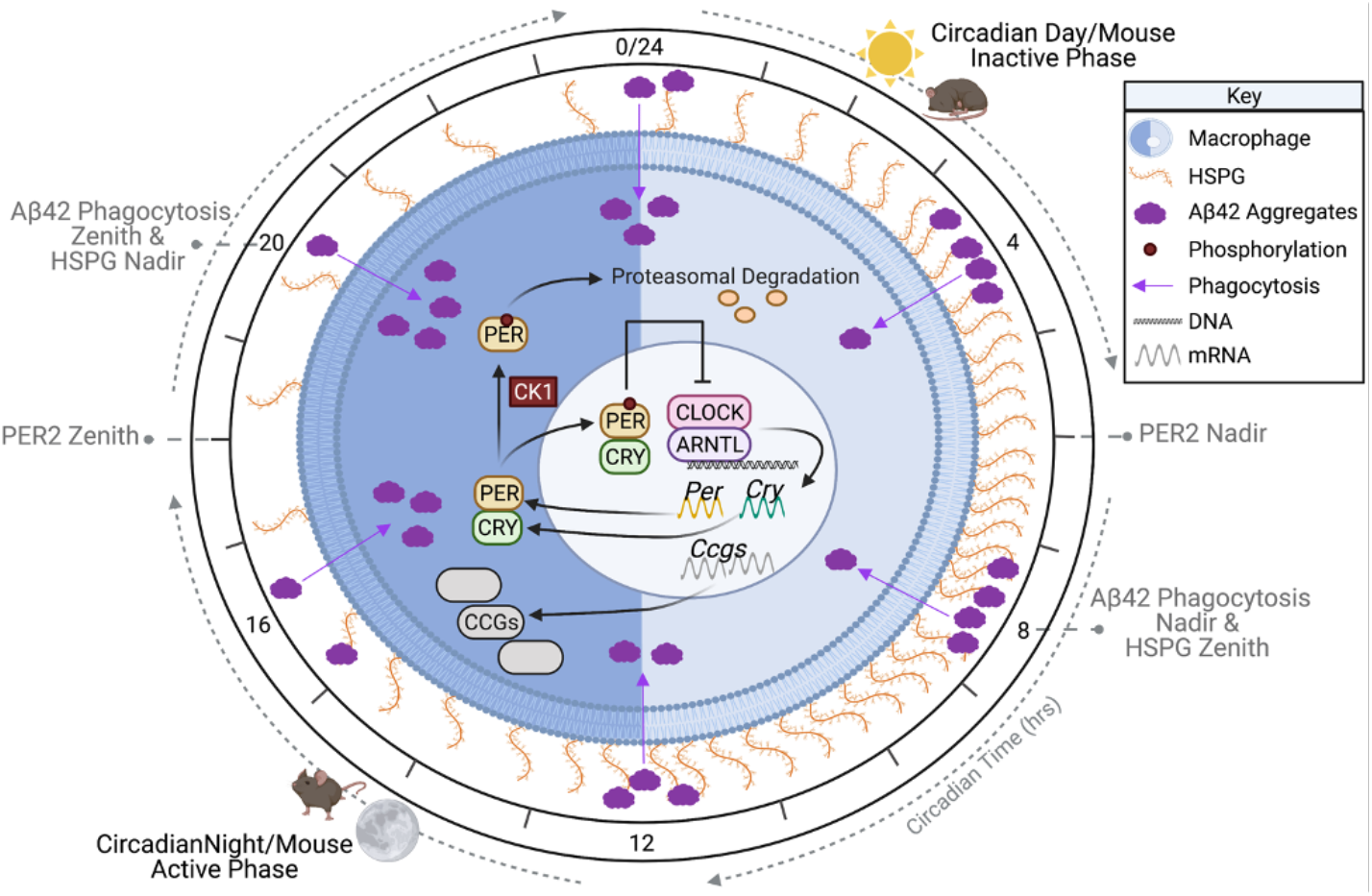

## Introduction

Alzheimer’s Disease (AD) and Alzheimer’s related dementias affect millions of people every year, are a leading cause of death in the U.S., and have associated care costs estimated at US$818 billion globally^1,2^. AD is a neurodegenerative, neuroinflammatory disease that is characterized by extracellular β-amyloid (Aβ) plaques, intracellular hyperphosphorylated tau fibrils, and increased neuroinflammation^3^. Though targeting Aβ as a therapeutic strategy has met limited success, Aβ accumulation is still regarded as a crucial step in AD pathogenesis due to strong evidence from human genetics of familial AD and Down syndrome^4,5^. Thus, understanding the metabolism of Aβ is essential to understand AD mechanisms and develop AD therapies.

A physiological consequence (and perhaps causative factor) of AD is the disruption of the circadian clock, the 24-hour endogenous rhythm that tunes physiology to the day/night cycle^6–10^ (Figure S1). Correlated with this, there is a daily oscillation in the abundance of Aβ42 in cerebrospinal fluid in healthy adults and this oscillation is ablated in patients with AD^6^. A key factor in the clearance of Aβ are the resident microglia and, in the later stages of AD, peripheral macrophages^7,11–13^. Increasing neuroinflammation due to the accumulation of Aβ42 plaques leads to elevated levels of macrophage markers, activating microglia and increasing peripheral macrophage migration across the blood brain barrier (BBB), where peripheral macrophages more efficiently clear Aβ42^7,11–17^. A double-edged sword, increased microglial activation and peripheral macrophage migration enhances the already high levels of neuroinflammation, leading to heightened cell death and exacerbating disease phenotypes^18–25^. The circadian clock also exerts extensive influence over macrophage/microglial behavior and disruption of circadian regulation affects the ability of macrophages to phagocytize target particles^26–28^ (Figure S1). In total, the concordance of these factors points to a relationship between the circadian regulation of the immune system and AD through the metabolism of Aβ42. Despite the correlation between circadian Aβ42 abundance and circadian control of macrophage/microglial phagocytosis, the link between AD and the clock via the circadian timing of Aβ42 phagocytosis has not been examined.

In this report, we utilized bone-marrow derived macrophages (BMDM’s) as a proxy for cells from the monocyte lineage to demonstrate that Aβ42 phagocytosis is under circadian control. Transcriptomic and proteomic data from macrophages identified few components that are circadianly regulated in the classical phagocytosis pathways but highlighted oscillations in the enzymes of the biosynthesis pathways of cell surface proteoglycans (PGs), which are known to negatively regulate Aβ42 phagocytosis^28–30^. We validated that PG levels oscillated over circadian time in macrophages *in vitro*, with PG levels reaching their zenith antiphase to the zenith of Aβ42 phagocytosis^29–35^. Chemically reducing PG levels ablated the circadian oscillation of Aβ42 phagocytosis by enhancing Aβ42 phagocytosis at its nadir. These data suggested that the presence of PGs suppresses the phagocytosis of Aβ42 and our investigation into the mechanism behind this suppression showed that aggregation and PG binding were essential to the circadian regulation of Aβ42 phagocytosis. Overall, our data suggests a role for myeloid cells in the circadian timing of the clearance of Aβ42 and an avenue through which the disruption of circadian rhythms can lead to enhanced AD pathogenesis.

## Results

### Phagocytosis of Aβ42 by Bone Marrow Derived Macrophages is Timed by the Circadian Clock

Aβ42 abundance oscillates with a circadian period, microglia and macrophages have been shown to phagocytize Aβ42, and phagocytosis by macrophages is under circadian regulation, leading us to hypothesize that oscillations in the metabolism of Aβ42 may stem from the circadian regulation of phagocytosis in cells from myeloid lineages^12,14,28^. To validate this theory, we first needed to demonstrate oscillations in the phagocytosis of Aβ42. To do so, we modified a previously-employed BMDM phagocytosis assay (as BMDMs are both models for activated microglia and are known to migrate into the brain in late-stage AD), to use fluorescently labeled Aβ42 to determine if the phagocytosis of Aβ42 is controlled by the circadian clock^28,36–38^. In brief, BMDMs were derived from bone marrow progenitor cells extracted from Per2::Luc C57BL/6J mice and differentiated with recombinant Macrophage Colony Stimulating Factor (M-CSF), with flow cytometry confirming complete differentiation into naïve peripheral macrophages^39,40^. These BMDMs were then serum-shock synchronized and luminescence traces confirmed that our protocol resulted in reliable, ∼24-h, PER2 oscillations^28^. 16 h after synchronization, confluent dishes of these BMDMs were treated HiLyte 488 labeled Aβ42 (Anaspec), in triplicate, every 4 h over a 24 h circadian period. BMDMs treated with labeled Aβ42 were harvested two hours after Aβ42 treatment and fixed with formalin. Total cellular fluorescence levels were analyzed with fluorescent confocal microscopy and quantified using a custom cell measurement MATLAB script (Figures 1A, S2, and Table S1).

**Figure 1.**
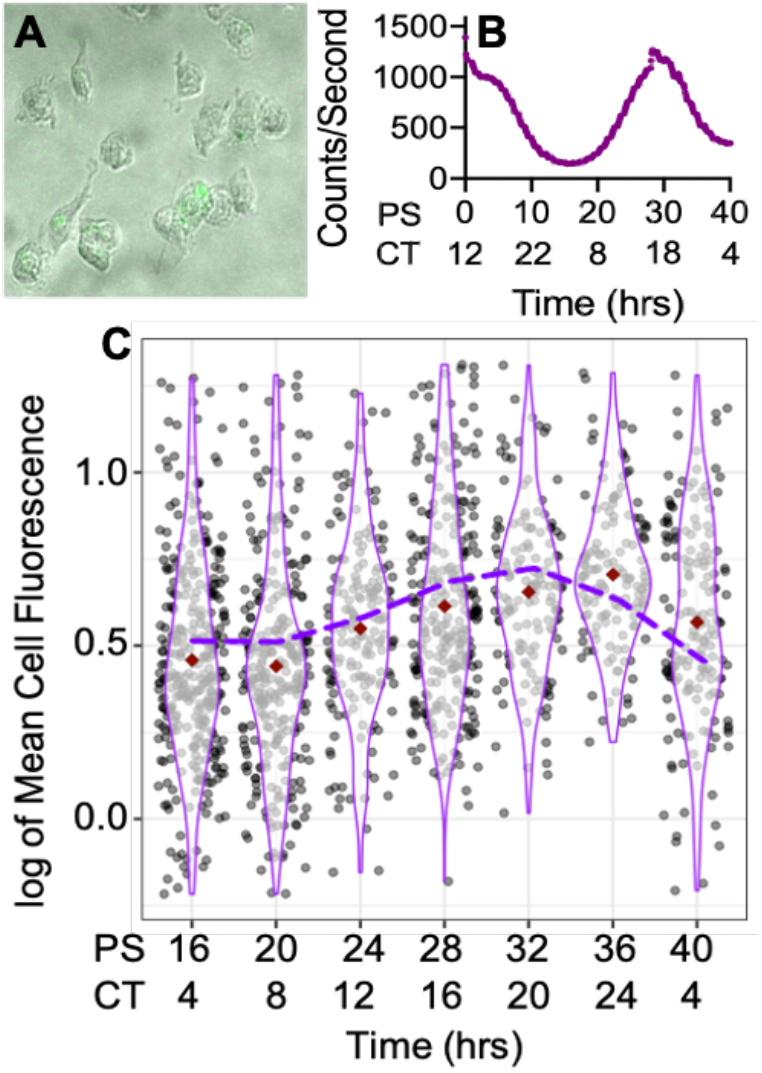
Aβ42 phagocytosis oscillated with a circadian period. (A) Cropped composite image of fluorescent microscopy of the Aβ42 treated macrophages. (B) PER2::LUC expression from untreated BMDMs, determined via bioluminescence using a lumicycle, reported in post shock (PS) and circadian time (CT). (C) Violin plot of Aβ42 phagocytosis represented in logged mean cell fluorescence values plotted against post shock (PS) and circadian time (CT). Grey dots represent a single cell measurement, purple violin plot lines represent the range of the data, and red diamonds indicate the mean measurement of each time point. The purple dashed line depicts the ECHO fitted model of Aβ42 phagocytosis. CT is in reference to the comparison of PER2 expression from Keller et al., 2009^41^ compared to PER2 expression from our cells.

Total cellular fluorescence levels demonstrated that the amount of Aβ42 phagocytized by macrophages underwent a daily oscillation, with the nadir occurring at 20 h post serum shock (PS) (CT8 in relation to the zenith of PER2-LUC) and the zenith at PS32 (CT20)^41^ (Figure 1B, 1C and S1). The extended harmonic circadian oscillator model (ECHO version 4.1) was used to analyze the average of the three replicates from each timepoint and predicted that Aβ42 total cellular florescence, and therefore the phagocytosis of Aβ42, oscillated with a circadian period (∼26 h, ECHO p-value= 1.61 × 10^−6^)^42^. Additionally, when compared with PER2 levels extrapolated based on the lumicycle data of the macrophages sampled, we saw that total cellular fluorescence was highest after the zenith of PER2 levels^40^ (Figure 1B and 1C). The Aβ42 phagocytosis experiment was then repeated using PER1^-/-^/PER2^-/-^ knockout (KO) mice to confirm the circadian influence of this relationship. BMDMs from the knockout mice were extracted, derived into naïve macrophages, and synchronized as described above. KO macrophages were treated with fluorescently labeled Aβ42 for 2 h, in triplicate, every 4 h for 24 h starting at PS16. Total cellular fluorescence was quantified and showed an ablation of the zenith of fluorescence at PS32 (CT20), confirming that the clock regulates Aβ42 phagocytosis (Figure S3).

### Proteoglycan Levels in Macrophages Undergo Daily Oscillations

We next used the ECHO program to probe our previously published, highly sampled/replicated, murine macrophage dataset, which tracked the transcriptome and proteome of BMDMs over two circadian days, to determine mechanisms that underlie the oscillation of Aβ42 phagocytosis^28,42^. Many known pathways essential for Aβ42 phagocytosis oscillated with a circadian period at the transcript level, but we found no oscillation in the proteome associated with these pathways except for the low-density lipoprotein receptor (LDLR) (zenith at PS25/CT13) and the low-density lipoprotein receptor-related protein isoform 6 (LRP6) (zenith at PS13/CT1)^43,44^ (Table S3). However, neither of these proteins reached their zenith in-phase with the oscillation of Aβ42 phagocytosis.

We then probed the two most enriched categories in the BMDM proteome according to Panther, binding and catalysis, and found several proteins related to proteoglycan (PG) synthesis and maintenance^45^. PGs are structurally diverse macromolecules that participate in a wide range of biological functions such as modulation of inflammation, extracellular matrix (ECM) assembly and remodeling, tissue repair, and ligand-receptor interactions^46,47^. PGs like heparan sulfate proteoglycans (HSPGs) and chondroitin sulfate proteoglycans (CSPGs) are involved in the clearance of Aβ42, associate with Aβ42 to foster the formation of plaques, and reduce Aβ plaque load upon the knockout of a key enzyme in the HSPG biosynthesis pathway^29–31,33–35,48–53^. Using the published PG synthesis pathway from Maeda, 2015^54^, the HSPG synthesis pathway from Kreuger and Kjellen, 2012^55^, and the CSPG synthesis pathway from Ly et al., 2011^56^, we cross-referenced the PG synthesis pathway enzymes to the mouse genome using Mouse Genome Informatics (MGI) to establish the homologues for PG synthesis in mice^57^. Once potential PG biosynthesizing genes were identified, we mapped each protein (or transcript if the protein was not found in our dataset) in these pathways to the information from our circadian transcriptomic and proteomics data sets and found that HSPG and CSPG biosynthesis enzymes undergo a daily oscillation at almost every step of PG biosynthesis in macrophages, from the formation of the linkage region to chain modification (Figure 2).

**Figure 2.**
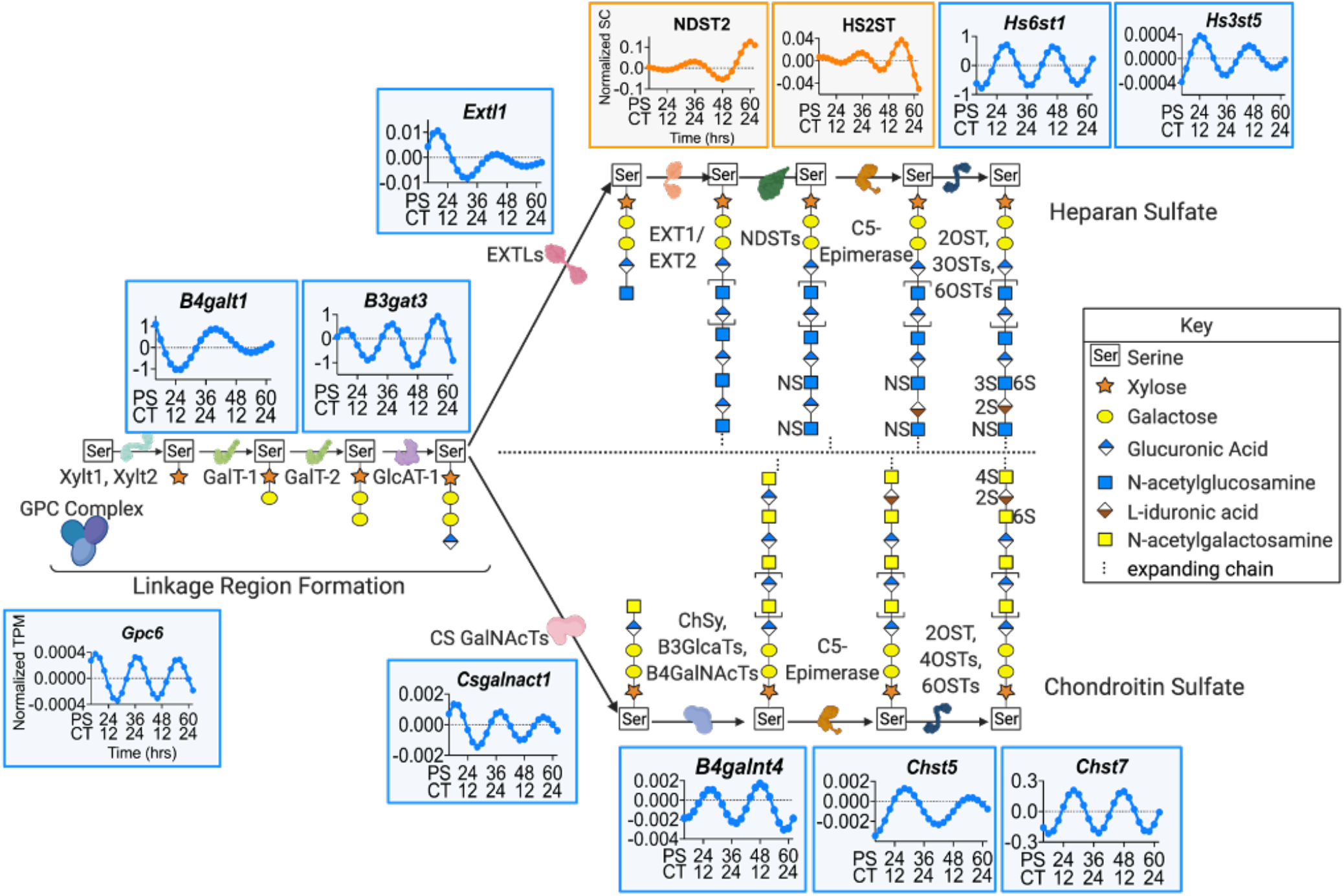
Enzyme levels in the HSPG and CSPG biosynthesis pathways varied over the circadian day. A diagrammatic representation of the pathways in HSPG and CSPG biosynthesis with the enzymes involved in each pathway that are under circadian regulation mapped to their location in the pathway. Pathways were adapted from Maeda, 2015^54^; Kreuger and Kjellen, 2012^55^; and Ly et al., 2011^56^. The plotted expression graphs depict the ECHO fitted model of the gene expression for either RNA (blue) or protein (orange) over PS and CT. Representative HSPG and CSPG chains are shown.

According to the current understanding of PG biosynthesis, creation of both HSPGs and CSPGs start with the formation of the linkage region by xylosyltransferases, XYLT1 and XYLT2, in conjunction with glypican proteins, GPC (*Gpc6* zenith at PS18.31/CT6.31). Galactosyltransferases, GALT-1 and GALT-2 (*B4galt1* zenith at PS40.35/CT8.35), sequentially facilitate the addition of two galactose residues, followed by the glucuronosyltransferase, GLCAT-1 (*Bgat3* zenith at PS19.1/CT 7.1), catalyzed addition of glucuronic acid. The HSPG and CSPG biosynthesis pathways then diverge and both branch-point enzymes exostosin-like glycosyltransferase, EXTLs (*Extl1* zenith at PS20.44/CT8.44), in the HSPG pathway and chondroitin sulfate N-acetylgalactosaminyltransferase, CS GALNACT (*Csgalnact1* zenith at PS19.01/CT7.01), in the CSPG pathway are under the regulation of the circadian clock. HSPG chains undergo elongation through the alternating additions of glucuronic acid and N-acetylglucosamine by exostosin glycosyltransferases, EXT1 and EXT2, modification by N-sulfotransferases, NDSTs (NDST2 zenith at PS34.61/CT2.61), the conversion of D-glucuronic acid into L-iduronic acid by C5-epimerase, and the addition of sulfate groups by sulfotransferases (*Hs6st1* zenith at PS37.53/CT5.53; *Hs3st5* zenith at PS24.72/CT16.72; *Hs2st* zenith at PS16.37/CT 4.37). In the CSPG pathway, CSPG elongation is catalyzed by chondroitin sulfate synthase, CHSY, along with glucuronosyl transferases, B3GLACTs, and N-acetyl galactosaminyl transferases, B4GALNACTs (*B4Galnt4* zenith at PS26.87/CT18.87). The chain is then modified by C5-Epimerase, and various sulfotransferases (*Chst5* zenith at PS28.71/CT7.71 and *Chst7* zenith at PS28.11/CT7.01) add sulfate groups (Figure 2 and Table S2). Clearly, the enzymes involved in the synthesis pathway for both HSPGs and CSPGs are tightly timed by the circadian clock.

To confirm that the oscillations we noted in PG synthesizing enzymes lead to oscillations of PG levels in macrophages, we extracted bone marrow from Per2::Luc mice and differentiated the extracted monocytes into BMDMs. We then grew these BMDMs to confluency, synchronized using serum shock, and harvested the BMDMs in quadruplicate every 4 h for 24 h, beginning at 16 h PS. Additionally, spent media and extracellular matrix (ECM) scrapings were sampled at four timepoints over 24 h (Figure S4A). These samples were then analyzed using Liquid Chromatography Tandem Mass-Spectrometry (LC-MS/MS) for HSPG and CSPG disaccharides in various sulfation states (Table 1). LC-MS/MS results were normalized by employing the CyQUANT cell proliferation assay at each time point to determine the total number of cells per pellet (Figure S4B).

**Table 1.**
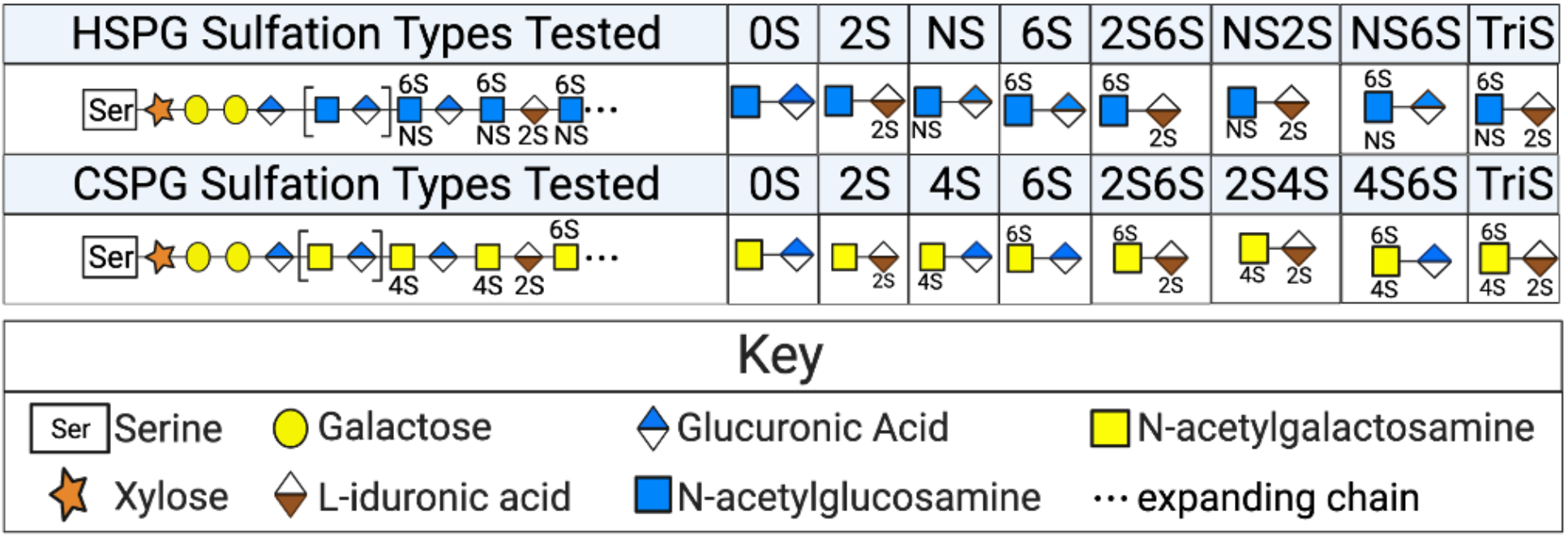
PG sulfation states quantified by LC-MS/MS. A schematic representation of the sulfation states tested in the cell samples, spent media, and ECM LC-MS/MS macrophage analysis. HSPGs investigated (top row) included 0S (no sulfation), 2S (sulfation on carbon 2), NS (sulfation on the amide), 6S (sulfation on carbon 6), 2S6S (sulfation on carbons 2 &6), NS2S (sulfation on the amide group and on carbon 2), NS6S (sulfation on the amide group and carbon 6), and TriS (sulfation on carbons 2 & 6 and on the amide group). CSPGs investigated (bottom row) include 0S (no sulfation), 2S (sulfation on carbon 2), 4S (sulfation on carbon 4), 6S (sulfation on carbon 6), 2S6S (sulfation on carbons 2 & 6), 2S4S (sulfation on carbon 2 & 4), 4S6S (sulfation on carbons 4 & 6), and TriS (sulfation on carbons 2, 4, and 6).

Our LC-MS/MS analysis of the macrophage cell pellet samples over circadian time demonstrated that, overall, the levels of both HSPGs and CSPGs oscillated with a circadian period (ECHO p-values =1.30 × 10^−7^ and 1.62 × 10^−7^ for HSPGs and CSPGs, respectively) (Figure 3A). Overall, circadian oscillations in the levels of PGs synthesis and expression reached their zenith at or near PS20/CT8. Interestingly, the oscillation of Aβ42 phagocytosis occurred antiphase to HSPG and CSPG expression, suggesting the presence of PGs inhibited Aβ42 phagocytosis (compare Figure 1 to Figure 3).

**Figure 3.**
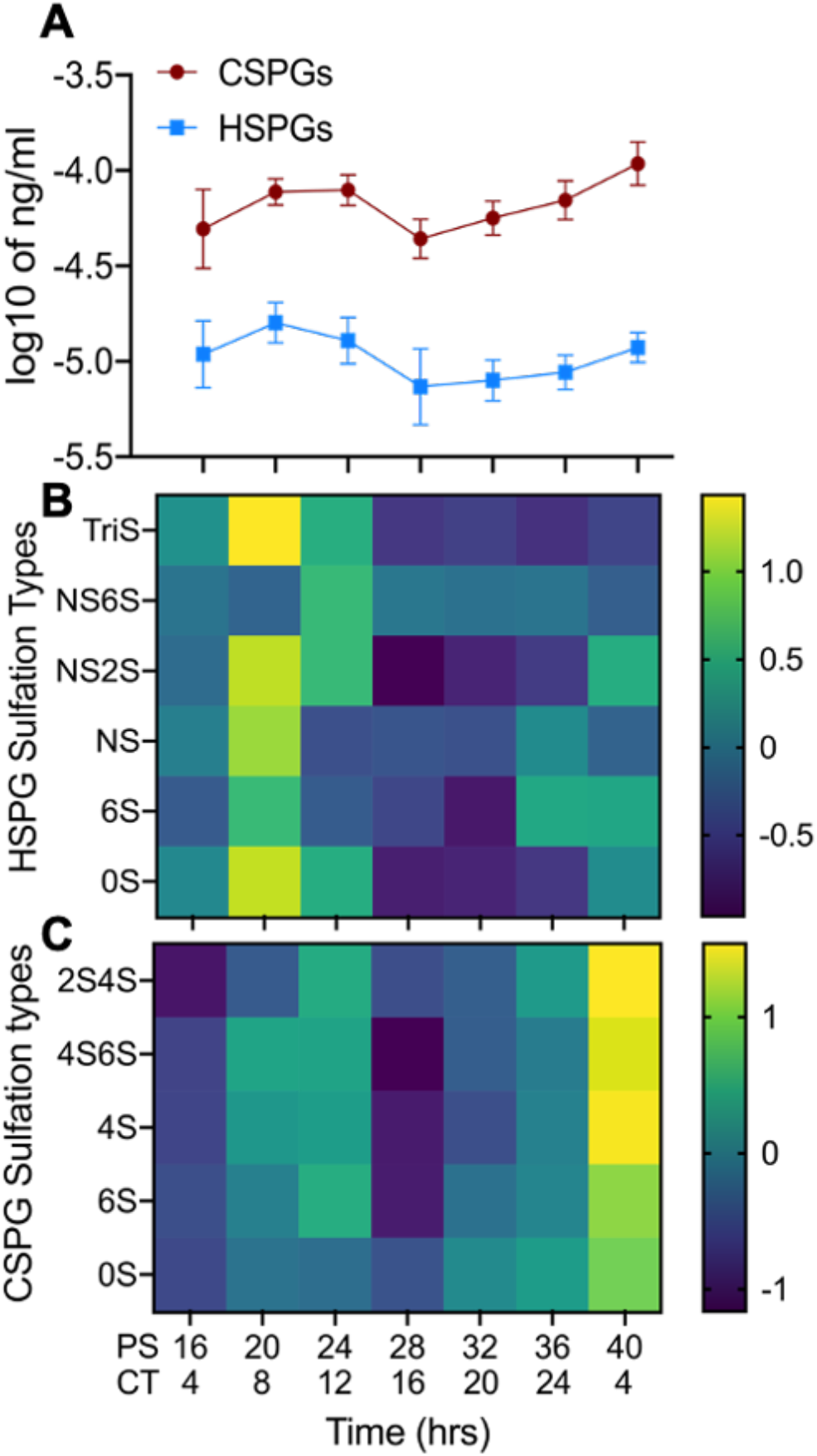
HSPG and CSPG levels in macrophages oscillated with a circadian period. (A) The sum of total cellular CSPG (n=4) and HSPG (n=4) levels (ng/cell) plotted against post shock (PS) and circadian time (CT). (B) Heat map of changes in cellular concentrations of specific HSPG sulfation types (n=4) shown in Z-scores over PS and CT time. (C) Heat map of changes in the concentrations of specific CSPG sulfation types (n=4) shown in Z-scores over PS and CT time. n = number of replicates, error bars represent the standard deviation of the replicates.

When all detected species of HSPGs identified in the cell pellet (0S, 6S, NS, NS2S, TriS, and NS6S) were analyzed individually, they were all found by ECHO to oscillate with a circadian period (Figure 3B, S5A, and Table S4). Similarly, when the detected species of CSPGs identified in the cell pellet (2S4S, 4S6S, 4S, 6S, and 0S) were analyzed individually by ECHO (Figure 3C, Figure S5D, and Table S4), they were all classified as oscillating with a circadian period. In the spent media samples, all detected HSPG and CSPG types (NS6S, NS2S, NS, and 0S for HSPGs and 0S, 4S, and 6S for CSPGs) were not found to be oscillating with a circadian period by ECHO (Figure S5C and F).

Finally, fewer HSPG and CSPG species were detected in the ECM scrapings (TriS, NS6S, NS2S, and 0S for HSPGs and 4S for CSPGs) and these species also appeared to have a daily oscillation (Figure S5B and E). We noted that in ECM samples the zenith in both HSPG and CSPG levels is at PS28 (Figure S5B and E). We also found in our proteomic data that proteins related to cell division such as cyclin dependent kinases, CDK1 and CDK2, CHK2 (a key regulator of cell cycle progress into mitosis), RAD21 (a cohesion complex component), and CDCA5 (*Sororin*, a chromatin-associated cohesion complex stabilizer), reached their zenith at PS28^28,58–60^ (Table S3). As HSPGs and CSPGs are known to be involved in ECM remodeling, an essential step in mitosis, this implies that the timing of the zenith of ECM PGs may represent an additional link between cell division and the circadian clock^46,61,62^.

### Rhythmic Phagocytosis of Aβ42 is Differentially Regulated by High-and Low-Sulfated HSPGs

We hypothesized that the rhythmic oscillation of HSPGs may inhibit the phagocytosis of Aβ42 due to the antiphase relationship between Aβ42 phagocytosis and PG levels and the published repressive effect of HSPGs on Aβ42 phagocytosis^29,30^ (Figures 1 and 3). To validate this hypothesis, we individually purified Heparinases I, II, and III, removed endotoxins, and confirmed Heparinase enzymatic activity (see methods)^63–66^ (Table S5). We then extracted, derived, synchronized, and exposed BMDMs to fluorescently-labeled Aβ42 every 4 h over one circadian day. In addition to Aβ42, we added a mixture of Heparinase I, II, and III to cleave all forms of HSPGs, with each biological replicate receiving a step-wise decreasing concentration of the mixture of the three heparinases (see Table S5). We found that regardless of Heparinase concentration, the addition of Heparinases I, II, and III ablated the oscillation in Aβ42 phagocytosis (Figure 4A). When comparing the relative levels of Aβ42 phagocytosis between the Heparinase treated and untreated macrophages, we found that Heparinase treatment increased phagocytosis when Aβ42 rhythmic phagocytosis reached its nadir (Heparinase Treatment PS20 vs Aβ42 PS20 Welch’s T-test p<2.2×10^−16^ and Hedges’ g=0.724791) but did not enhance overall levels of phagocytosis as compared to Aβ42-alone due to the low Hedges g value (Heparinase treatment PS32 vs Aβ42 PS32 Welch’s T-test p=0.006426 and Hedges’ g=0.361095) (Figure 4). This data confirmed that circadianly-timed increases in the levels of HSPGs were rhythmically inhibiting the phagocytosis of Aβ42.

**Figure 4:**
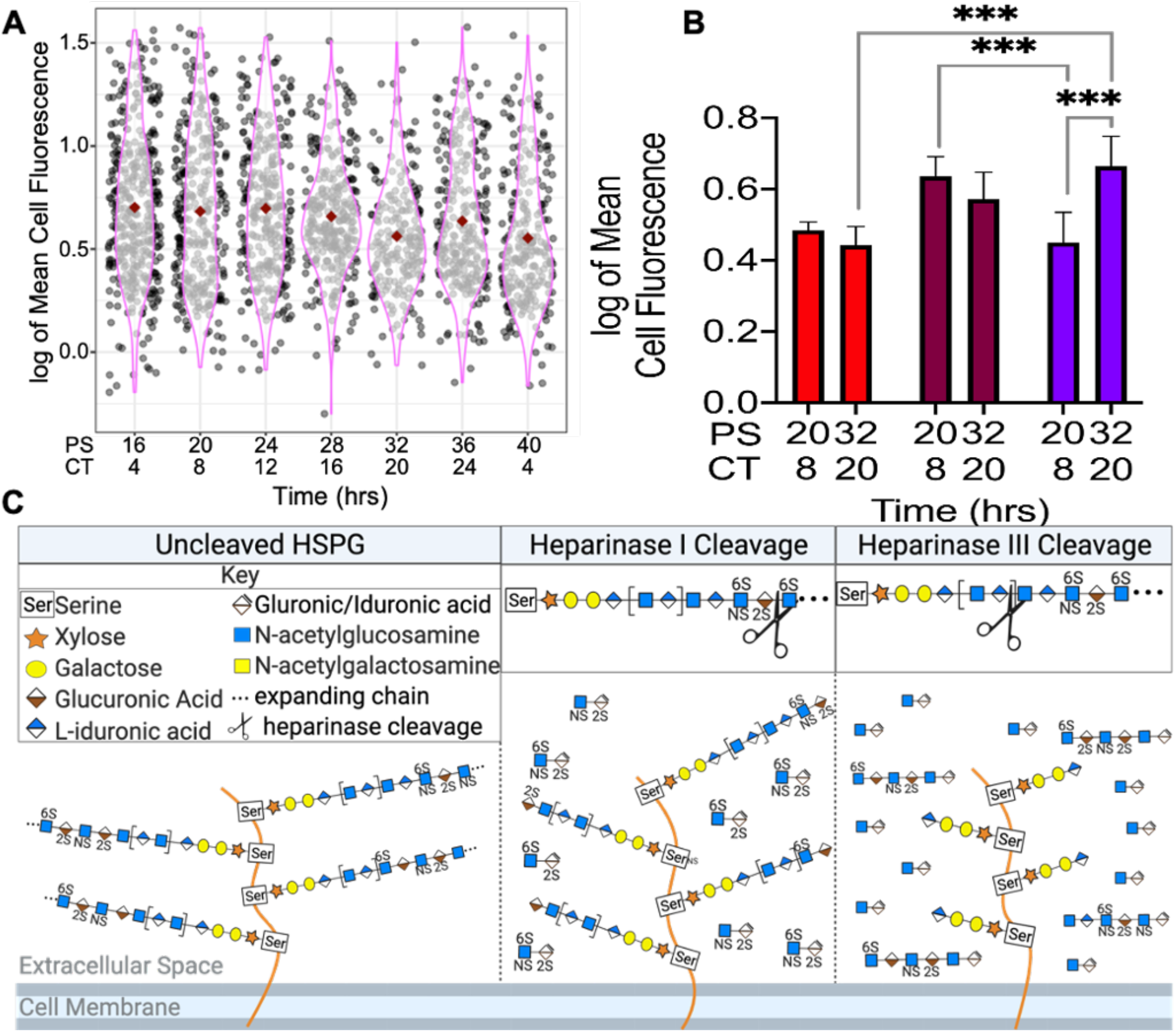
Heparinase treatment ablated daily oscillations in Aβ42 phagocytosis in macrophages. (A) A violin plot showing the phagocytosis of Aβ42 in the presence of a mixture of heparinases I, II, and III in logged mean fluorescence values plotted against PS and CT. Each cell sampled per time point is represented by a black or gray dot with the mean shown as a red diamond. The pink outline represents the range of values per timepoint. (B) A bar graph comparing Aβ42 phagocytosis when heparinases I or III were added at PS20 (CT8) and PS32 (CT20). Red bars denote Heparinase 1 + Aβ42, maroon bars denote Heparinase 3 + Aβ42, purple bars denote Aβ42 without heparinases. All data shown was performed in triplicate and the error bars represent the standard deviation of the replicates. ***= p < 2.2 × 10^−16^ and Hedge’s G > 0.5 (C) Images depicting where individual heparinase cleave representative HSPG chains.

Notably, our data demonstrated that 0S HSPG was the predominant HSPG species in macrophages, indicating that 0S HSPG may be the principal factor that leads to the circadian inhibition of Aβ42 phagocytosis. To determine if our theory was correct, we next analyzed the phagocytosis of Aβ42 at the zenith and nadir of Aβ42 phagocytosis (PS20 and PS32) in the presence of either Heparinase III or Heparinase I only (based on Figure 3 and Table S4). Isolated BMDMs, at PS20 and PS32 were treated in triplicate with HiLyte-488 labeled Aβ42 and actively-equal amounts of either Heparinase I or Heparinase III and analyzed as described above (Table S5). Heparinase I specifically cleaves between L-iduronic acid and N-acetylglucosamine on highly sulfated chains in HSPGs, leaving lower sulfated chains bound to the cell surface. Heparinase III specifically cleaves between glucuronic acid and N-acetylglucosamine on lower-sulfated chains in HSPGs, leaving fragments of highly-sulfated heparan sulfate chains in the solution (Figure 4C).

We found that in macrophages that had been treated with Heparinase I, there was a loss of daily oscillation compared to untreated (Aβ42-Hep 1 PS20 vs PS32 Welch’s T-test p= 0.08295 and Hedges’ g= 0.101721; Aβ42-alone PS20 vs PS32 Welch’s T-test p= 8.979 × 10^−15^ and Hedges’ g= 0.764648) and overall levels of phagocytosis significantly decreased at the circadian zenith, but not the nadir, as compared to Aβ42 phagocytosis in untreated macrophages (zenith: Aβ42-Hep 1 PS32 vs Aβ42-alone PS32 Welch’s T-test p=2.2 × 10^−16^ and Hedges’ g=0.635008; nadir: Aβ42-Hep 1 PS20 vs Aβ42-alone PS20 Welch’s T-test p=0.06002 and Hedges’ g=0.132791) (Figure 4 and S6A). When macrophages were treated with Heparinase III, oscillations in Aβ42 phagocytosis were also ablated as compared to untreated (Aβ42-Hep 3 PS20 vs PS32 Welch’s T-test p=0.001997 and Hedges’ g=0.1663366). Conversely, phagocytosis increased at the nadir but not the zenith, as compared to Aβ42 phagocytosis in untreated macrophages (nadir: Aβ42-Hep 3 PS20 vs Aβ42-alone PS20 Welch’s T-test p<2.2 × 10^−16^ and Hedges g=0.5408882; zenith: Aβ42-Hep 3 PS32 vs Aβ42-alone PS32 Welch’s T-test p= 0.0007397 and Hedges’ g=0.21997) (Figure 4 and S6B). The demonstration that the absence of high or low sulfated species of HSPGs on the cell surface differentially ablated circadian oscillations in Aβ42 phagocytosis indicated that the proper ratio of cell-surface sulfation states of HSPGs regulated the circadian timing of Aβ42 phagocytosis.

### Binding of HSPGs to Aβ Peptides is a Repressive Factor in the Circadian Phagocytosis of Aβ42

Highly sulfated HSPGs, like perlecan, and CSPGs, like chondroitin-4S and dermatan sulfate, have been shown to directly interact with Aβ^50,52,67^. The interaction between Aβ42 and heparan sulfate is known to occur through electrostatics from the dense negative charge of heparan sulfate and positively charged residues of Aβ42, particularly the HHQK cluster on Aβ42 (amino acids 13-16)^68–71^. Mouse Aβ42 (mAβ42) is nearly identical to human Aβ42, with only three amino acid substitutions (R5G, Y10F, and H13R). However, these differences, particularly the substitution of an arginine at the histidine residue, reside in or near the HHQK cluster and affects the interaction between HSPGs and mAβ42 as compared to human Aβ42^71–73^ (Figure 5C). This difference is important as while mice naturally produce mAβ42, mAβ42 plaques do not form and there is no natural occurrence of AD in mice, suggesting binding between HSPGs and Aβ42 may play a role in the development of AD disease phenotypes^74–76^.

**Figure 5:**
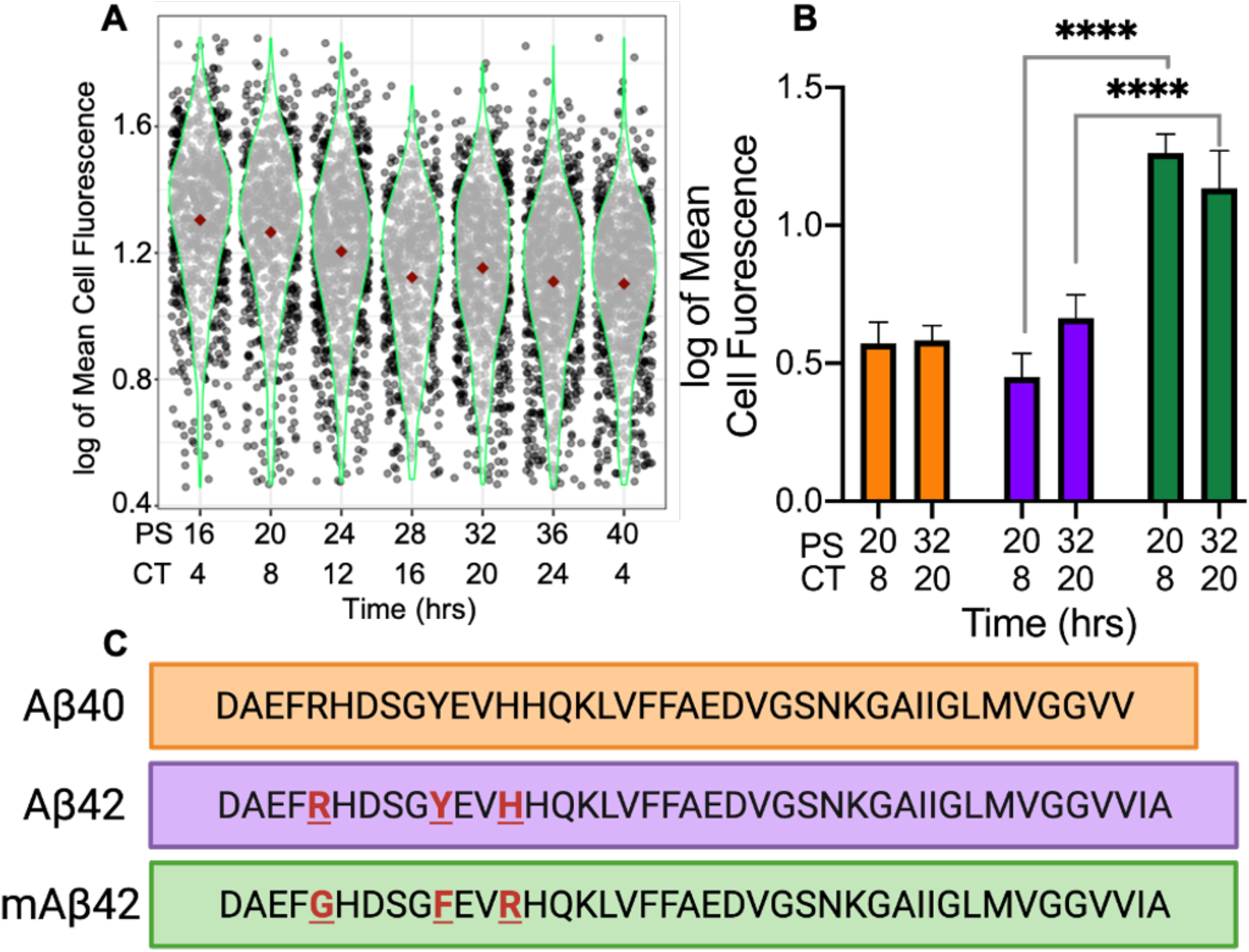
HSPG-Regulated Circadian Macrophage Phagocytosis was Contingent upon HSPG-Aβ Binding and Aggregation. (A) Violin plot of mAβ42 phagocytosis represented in logged mean cell fluorescence values plotted against PS and CT. Grey dots represent a single cell measurement, green violin plot lines represent the range of the data, and red diamonds indicate the mean measurement of each time point. (B) A bar chart comparing Aβ40 phagocytosis (orange) to Aβ42 phagocytosis (purple) and mAβ42 phagocytosis (green) at PS20/CT8 and PS32/CT20. All data shown was performed in triplicate and the error bars represent the standard deviation of the replicates. ****= p<2.2 × 10^−16^ Hedge’s g>2.0 (C) An amino acid sequence comparison to show the differences between the Aβ42, Aβ40, and mAβ42 sequences.

This led us to hypothesize that binding between HSPGs and Aβ42 at the cell surface may play a mechanistic role in the circadian phagocytosis of Aβ42. We therefore repeated our phagocytosis experiment using mouse HiLyte 488 labeled mAβ42 (Anaspec). Bone marrow derived from Per2::Luc mice was used to generate synchronized macrophages and, in triplicate, mAβ42 phagocytosis was assayed an analyzed as described above every 4 h for 24 h. mAβ42 phagocytosis did not have a circadian oscillation according to ECHO, suggesting that HSPG binding is essential for HSPG to impart circadian timing of Aβ42 phagocytosis in macrophages^42^ (Figure 5A). Aligned with our hypothesis, and with the data suggesting that HSPGs inhibit the phagocytosis of human but not mouse Aβ42, the resulting data showed a significant increase in mAβ42 phagocytosis at the zenith and nadir of human Aβ42 uptake (mAβ42 PS20 vs Aβ42 PS20 Welches T test p<2.2 × 10^−16^, Hedges’ g=3.031588; mAβ42 PS32 vs Aβ42 PS32 Welch’s T test p<2.2 × 10^−16^, Hedges g=2.020813) as well as increased phagocytosis compared to human Aβ42 phagocytosis at all sampled times^30^ (Figure 5B). This data, correlated with the zenith in HSPG levels occurring at the nadir in Aβ42 phagocytosis (Figures 1 and 3), suggested the circadian timing of increased cell-surface HSPG levels, and therefore interactions, negatively regulates the phagocytosis of Aβ42.

### Circadian Phagocytosis of Aβ42 is Correlated with Aβ42 Plaque Formation

In addition to binding Aβ42, HSPGs regulate the formation of Aβ42 plaques, with highly-sulfated HSPGs accumulating in the plaques^50,52,67,72,77^. Aβ peptides exist in two forms (Aβ40 or Aβ42) based on the cleavage of the amyloid precursor protein (APP) by β-and γ-secretase^78^. Previous studies show differential phagocytosis of Aβ42 and Aβ40, yet the key amino acids essential to binding are present in Aβ40 (Figure 5C) (13-16, HHQK)^29,79^. However, the anionic bridge between lysine 28 and alanine 42, which is broken by HSPGs to allow the formation of Aβ42 aggregates, does not occur in Aβ40, and thus HSPGs do not affect Aβ40 aggregation^71,80^. Correlated with this, the Aβ42 peptide shows higher aggregation kinetics and higher toxicity than Aβ40^81^.

Paired with our data showing an increase in phagocytosis when free highly-sulfated HSPG fragments are present (Figure 4), we hypothesized that if Aβ42 aggregation was a key factor in HSPG timing of circadian phagocytosis, then there should be no oscillation in the phagocytosis of Aβ40. We utilized our above-described phagocytosis assay to analyze the phagocytosis of Aβ40 at the zenith and nadir times of Aβ42 phagocytosis (PS20 and PS32) to determine if the there was an oscillation in the phagocytosis of Aβ40. We found no difference in Aβ40 phagocytosis at PS20 and PS32 (Aβ40 PS20 vs PS32 Welch’s T-test p=0.2362 and Hedges’ g=0.067337) confirming that there is no oscillation in Aβ40 phagocytosis, and when we compared total phagocytosis of Aβ40 at PS20 and PS32 to total phagocytosis of Aβ42 at PS20 and PS32, we found no difference in overall phagocytosis due to the low Hedges’ g value (Aβ40 vs Aβ42 Welch’s T-test p=1.278 × 10^−05^ and Hedges’ g=0.275235). Overall, our results confirmed our hypothesis that the HSPG-regulated circadian-timing of Aβ42 phagocytosis is also dependent upon Aβ42 plaque formation (Figure 5B).

## Discussion

The clearance of Aβ42, which is inhibited in AD pathology, is essential for a healthy neuronal microenvironment^6,82^. Metabolism of Aβ42 is controlled by the circadian clock *in vivo* and disruption of oscillations in this rhythm could affect the accumulation of Aβ42^6^. Accumulation of Aβ42 accelerates the development of the symptoms associated with AD, providing a possible mechanism for the positive correlation between AD and clock disruption^8,9^. To add to this connection, our results demonstrate circadian control of the phagocytosis of Aβ42 in murine macrophages (Figure 1). As peripheral macrophages are models for microglia and also migrate to the brain in the later stages of AD, our findings suggest the disruption of the circadian timing of macrophage/microglia phagocytosis may be a vital component in Aβ42 metabolism, highlighting a potential causative factor in the increase in accumulation of Aβ42 in patients with clock/sleep disturbances^8,13,20,22,25,82,83^.

Our analysis showed that a key factor in the circadian timing of Aβ42 phagocytosis was the presence of cell-surface HSPGs. LC-MS/MS analysis of macrophages over circadian time showed a distinct rhythm in HSPG levels (Figures 2 and 3). The zenith and nadirs of the HSPG oscillation were antiphase to those of Aβ42 phagocytosis, suggesting that HSPGs play an inhibitory role in the phagocytosis of Aβ42. The removal of all HSPG chains by treating synchronized macrophages with Heparinase I, II and III ablated the rhythm in the oscillation of Aβ42 phagocytosis by increasing phagocytosis at the nadir, cementing the importance of HSPGs in the circadian timing of Aβ42 clearance (Figure 4). Previous studies have shown differential effects of HSPG species on Aβ42, and our data paralleled this as we found that it is the ratio of HSPGs that is essential for the circadian timing of phagocytosis, though the complexity of the resultant HSPG fragments after individual Heparinase treatment make determining these roles difficult^48,49,52,84–86^ (Figure 4). While HSPGs have previously been implicated in multiple pathways of Aβ clearance, cellular toxicity, and plaque generation, our work is the first to demonstrate a link between the circadian timing of HSPGs and Aβ42 clearance^50,52,86–89^.

The relationship between Aβ42 clearance and HSPG binding has been shown, with HSPGs, perlecan, and syndecans binding Aβ42, regulating Aβ42 toxicity^51,72,88,90^. Highly sulfated HSPGs bind Aβ42 both extracellularly or at the cell surface to affect phagocytosis^67,86,91^. We predict that the circadianly-timed increase in HSPGs on the cell surface could affect HSPG/Aβ42 binding and this binding could be the mechanistic cause of the temporal inhibition in Aβ42 phagocytosis. Suppression of phagocytosis may be key to preventing plaque formation as a recent land-mark study found that microglial phagocytosis of Aβ and sequestration in lysosomes promotes plaque formation^92^ (Figure 1). Our data showing that the overall phagocytosis of mAβ42, which is predicted to not bind HSPGs as tightly as Aβ42, was higher and had no oscillation compared to human Aβ42, supports this theory^71–73^ (Figure 5).

Congruently, Aβ42 plaque formation has been positively correlated to the presence of HSPGs^49,67,86^. Previous studies have shown that low-sulfated HSPGs did not enhance Aβ42 plaque formation and lack of *N*-sulfated heparin reduced Aβ phagocytosis^52^. Our work correlated with this, as the phagocytosis of Aβ40, which forms far fewer plaques than Aβ42, has no circadian oscillation in phagocytosis, suggesting that plaque formation is also correlated with circadian phagocytosis of Aβ42^78^ (Figure 5). Aβ40 had similar levels of overall phagocytosis as compared to Aβ42, though this is possibly due to solubility differences as microglia readily clear soluble Aβ, or due to energy independent endocytosis of Aβ40, which has been demonstrated in neurons^93–96^. Moreover, when we released HSPGs from the cell surface using Heparinase III, making long sulfated HSPG chains readily available for the formation of aggregates, we saw an increase in phagocytosis at the nadir, supporting this hypothesis^77^ (Figure 4). Furthermore, the source of HSPGs involved in plaque formation is hypothesized to be from microglia and macrophages, providing a further potential connection between AD, the circadian clock, and the immune system^97–99^.

Beyond the discovery of the circadian timing of Aβ42 phagocytosis by HSPGs, there is a striking amount of circadian regulation devoted to timing oscillations in the levels of all PGs (Figure 3). We found that the circadian clock tightly controlled the pathways that regulate the production of HSPGs and CSPGs (Figure 2). As PGs are key regulators for cell surface interactions in the inflammatory response, the circadian regulation of PGs in macrophages has strong implications for inflammation and immunity beyond its affect in AD^100^. Macrophage polarization, which is also controlled by the circadian clock, results in differently expressed HSPGs, with less 2S and 6S sulfation when pushed into the M1 or inflammatory state^97,101,102^. This implies that the circadian regulation of HSPG and CSPG expression could be critical to preventing a hyper inflamed state in inflammatory diseases such as AD. Moreover, as PGs play critical roles in development and cellular processes, it is likely that PGs fall under the regulation of the circadian clock in other tissue types, making it a promising target for future circadian studies^100,103,104^.

## Acknowledgements

We would like to acknowledge Dr. Emily Collins for her transcriptome and proteome datasets, Lufeng Yu for performing heparinase activity assays, Ke Xia for providing editing on the LC-MS/MS section, and the PER1^-/-^/PER2^-/-^ knockout mice shared by the labs of Jay Dunlap and Jennifer Loros. Additionally, we would like to acknowledge the Analytical Biochemistry Core at RPI, Sergey Pryshchep of the Microscopy core at RPI for his training and guidance, and Dr. Antigone McKenna and the BioResearch core staff at RPI for their excellent care of our mouse lines. We would like to thank BioRender.com for support in figure creation. Finally, this work was supported by an NIH-National Institute of Biomedical Imaging and Bioengineering Grant U01EB022546 (to J.M.H), an NIH-National Institute of General Medical Sciences Grant R35GM128687 (to J.M.H.), an NSF CAREER Award 2045674 (to J.M.H.), NIH grants 1RF1AG069039 (to C.W.), DK111958 and CA231074 (to R.J.L.), Rensselaer Polytechnic Startup funds (to J.M.H.), a gift from the Warren Alpert Foundation (to J.M.H.), and a NIH-National Institute of Aging T32 Fellowship AG057464 (to G.T.C.).

## Author Contributions

G.T.C. wrote the manuscript, designed and carried out experiments, and analyzed results. Y.Y. carried out experiments and analyzed the resultant data. C.A.U. analyzed data and aided in the writing of the manuscript. G.F. provided data analysis. F.Z. planned experiments. R.J.L. and C.W. planned experiments, analyzed data, and edited the manuscript. J.M.H. planned experiments, analyzed data, and wrote the manuscript.

## Methods

### Animals

PER2::LUC C57BL/6J male mice 3-6 months of age bred from Jackson Labs stock strains (Accession # 006852) were used for all WT experiments and were euthanized using CO_2_ gas and cervical dislocation^40^. Per2::Luc C57BL/6J mice were kept on a strict lighting schedule of 12L:12D to maintain synchronized circadian rhythms and fed standard rodent chow *ad libitum*. PER1^-/-^/PER2^-/-^ knockout male mice 3-6 months in age were used in all KO experiments and were euthanized using CO_2_ gas and cervical dislocation^105^. PER1^-/-^/PER2^-/-^ mice were kept on a strict 12L:12D lighting schedule and were fed antibiotic rodent chow *ad libitum* due to a *Helicobacter* infection in this mouse line.

### Reagents

Amyloid-beta 42 labeled with HiLyte 488 (AS-60479-01), Amyloid-beta 40 labeled with HiLyte 488 (AS-60491-01), and custom Mouse amyloid-beta 42 labeled with HiLyte 488 (SQ-ANCPXXXX) were all purchased from Anaspec (Fremont, CA, USA). ACK lysing buffer (10-548E) was manufactured by Lonza.

### Media

For macrophage differentiation media, DMEM media (Corning 10-013-CV) was used and supplemented with fetal bovine serum (FBS) (Gibco 10437028) at a 1:10 dilution, 100x Penstrep at a 1:100 dilution (Corning MT30002CI), 200mM L-glutamine at a 1:100 dilution (Corning 25-005-CI), Beta-mercaptoethanol (BME) at a 1:1000 dilution (Gibco 21985-023), and 50 mg/ml Gentamicin at a 1:1000 dilution (Gibco 15750-060).

For macrophage assay medium (after serum shock), Leibovitz media (Gibco 21083-027) was used and supplemented with fetal bovine serum (FBS) at a 1:10 dilution (Gibco 10437028), 100x Penstrep at a 1:100 dilution (Corning MT30002CI), 200mM L-glutamine at a 1:100 dilution (Corning 25-005-CI), Beta-mercaptoethanol (BME) at a 1:1000 dilution (Gibco 21985-023), 1 M HEPES at a 1:100 dilution (Gibco 15630-080), 50 mg/ml Gentamicin at a 1:1000 dilution (Gibco 15750-060), and Luciferin at a 1:100 ratio supplied by (Gold Biotechnology LUCK-1G) when appropriate.

For macrophage serum starvation assay media, Leibovitz media (Gibco 21083-027) was used and supplemented with 100x Penstrep at a 1:100 dilution (Corning MT30002CI), 200 mM L-glutamine at a 1:100 dilution (Corning 25-005-CI), Beta-mercaptoethanol (BME) at a 1:1000 dilution (Gibco 21985-023), 1 M HEPES at a 1:100 dilution (Gibco 15630-080), and 50 mg/ml Gentamicin at a 1:1000 dilution (Gibco 15750-060).

For macrophage serum shock media, Leibovitz media (Gibco 21083-027) was used and supplemented with FBS at a 50% dilution (Gibco 10437028), 100x Penstrep at a 1:100 dilution (Corning MT30002CI), 200 mM L-glutamine at a 1:100 dilution (Corning 25-005-CI), Beta-mercaptoethanol (BME) at a 1:1000 dilution (Gibco 21985-023), 1M HEPES at a 1:100 dilution (Gibco 15630-080), and 50mg/ml Gentamicin at a 1:1000 dilution (Gibco 15750-060).

All cell culture medias contained Macrophage Colony Stimulating Factor (M-CSF) at a 1:1000 dilution (Prospec cyt-439-c). Phosphate buffered saline (PBS) (Corning 21-040-CV) was used to wash the cells in between media changes. Cell Stripper (Corning 25-056-CI) was used to remove the adherent macrophages from cell culture dishes for harvesting. Buffered formalin phosphate (Fisher Chemical SFL004) was used for fixing harvested cells.

### Mass spectrometry standards

Unsaturated disaccharide standards of CS (0S_CS-0_: ΔUA-GalNAc; 4S_CS-A_: ΔUA-GalNAc4S; 6S_CS-C_: ΔUA-GalNAc6S; 2S_CS_: ΔUA2S-GalNAc; 2S4S_CS-B_: ΔUA2S-GalNAc4S; 2S6S_CS-D_: ΔUA2S-GalNAc6S; 4S6S_CS-E_: ΔUA-GalNAc4S6S; TriS_CS_: ΔUA2S-GalNAc4S6S), (Iduron, CD001, CD002, CD003, CD004, CD005, CD006, CD007 and CD008, respectively), unsaturated disaccharide standards of HS (0S_HS_: ΔUA-GlcNAc; NS_HS_: ΔUA-GlcNS; 6S_HS_: ΔUA-GlcNAc6S; 2S_HS_: ΔUA2S-GlcNAc; 2SNS_HS_: ΔUA2S-GlcNS; NS6S_HS_: ΔUA-GlcNS6S; 2S6S_HS_: ΔUA2S-GlcNAc6S; TriS_HS_: ΔUA2S-GlcNS6S), and unsaturated 1,3-linked disaccharide standard of HA (0S_HA_: ΔUA-GlcNAc), (Iduron, HD006, HD005, HD008, HD007, HD002, HD004, HD003, HD001 and HA02, respectively) were purchased from Iduron, UK, where ΔUA is 4-deoxy-α-L-*threo*-hex-4-enopyranosyluronic acid.

### Enzymes

Chondroitin lyase ABC from *Proteus vulgaris* was expressed in the Linhardt laboratory (see below). Recombinant *Flavobacterial* heparin lyases I, II, and III were expressed in the Linhardt laboratory using *Escherichia coli* strains provided by Jian Liu (College of Pharmacy, University of North Carolina).

### Chemicals

2-Aminoacridone (AMAC) (Sigma-Aldrich 06627) and sodium cyanoborohydride (NaCNBH_3_) (Sigma-Aldrich 156159) Calcium chloride (449709), DMSO (Sigma-Aldrich 94563), acetic acid (Sigma-Aldrich A6283), ammonium acetate (Sigma-Aldrich 73594) and methanol (Sigma-Aldrich 646377) were obtained from Sigma-Aldrich (St. Louis, MO, USA).

### Bone Marrow Derived Macrophage Extraction and Synchronization

Bone marrow was harvested from the femurs and tibias of one 3–6-month-old male mouse using a 26 g needle and syringe filled with DMEM supplemented media. ACK lysing buffer (Lonza 10-548E) was used to lyse red blood cells in order to prevent erythrocyte contamination. Filtered bone marrow cells were counted using a BioRad TC20 automated cell counter and plated on 35 mm cell culture plates at a density of 1 × 10^6^ in macrophage differentiation media. After three days of incubation at 37°C with 5% CO_2_, fresh macrophage differentiation media was added. After another 3 days of incubation, the cells were washed with 2 ml warm PBS per plate, and replaced with macrophage assay media, and the cultures were incubated for 24 h. Macrophage cultures were then washed with 2 ml warm PBS, macrophage serum starvation media was added, and the plates were incubated in the starve media for 24 h. Next, macrophage serum starvation media was removed, and macrophage serum shock media was added, and cultures were incubated for 2 h^106^. Macrophage cultures were then washed with 2 ml warmed PBS each and macrophage media Luciferin was added. Cultures were sealed with grease and glass cover slips and placed in a LumiCycle 32 luminometer (Actimetrics) to confirm synchronization of circadian rhythmicity. Assays began 16 h post serum shock to allow the synchronized cells to return to homeostasis.

### Amyloid-beta Reconstitution

All amyloid-beta isoforms were reconstituted 24 h or less before experimentation in water and PBS (1:1) and were dissolved using sonication for approximately 30 min. The reconstituted Aβ was pooled from individual vials before aliquoting for each experimental replicate. The aliquots were then flash frozen and stored at -80°C until 20 min before use.

### Phagocytosis Assay

Using synchronized macrophages (as described above) starting at PS16 macrophages were treated every 4 h for 24 h, in triplicate, with 115 μl (0.25 mg/ml) of fluorescently labeled amyloid-beta (PS16, PS20, PS24, PS28, PS32, PS36, and PS40) and heparinases at the reported activity levels (Table S5). The treated macrophages were then incubated at 37°C for 2 h. Following the two-hour incubation, the media was aspirated, and the macrophages were washed with 2ml of PBS three times before Cell Stripper (Corning 25-056-CI) was added to the cultures, which were then incubated at 37°C for 3 min. Detached macrophages were next centrifuged at 400 *g* for 5 min, the supernatant was aspirated, and the macrophage cell pellets were resuspended in formalin and incubated in the dark at room temperature for 30 min. Macrophages were then centrifuged for 5 min at 400 *g*, the supernatant was aspirated, and the macrophages were resuspended in 250 µL of PBS and kept at 4°C. The fixed macrophages were imaged using fluorescent microscopy within a week of the experiment.

### Fluorescent Image Analysis

Macrophage cells containing fluorescent Aβ were imaged on a Zeiss LSM 510 Laser scanning confocal microscope using the 40x objective and an argon laser at 488nm excitation wavelength. Macrophages were viewed in two channels, the light channel and the fluorescent channel and microscope settings were saved and reused for consistency between experiments. Images were taken until 100+ macrophages were sampled. To analyze these images, we used a custom MATLAB script. This script uses defined cell size ratios and edge detection to pick out single cells and objects (up to 4 cells clumped together) and measures the average pixel intensity, max pixel intensity, and area of each object. This script thresholds out the background and subtracts this measurement from the average cellular pixel intensities. *Image analysis script will be publicly available on GitHub upon publication*.

### Statistical Analysis of Cellular Pixel Intensities

Average cellular pixel intensities were used as a proxy for of Aβ phagocytosis. At least 100 cells measured per time point were analyzed using the above-described software. All measurements below the background signal were considered below the threshold and removed. The average pixel intensities were then log10 transformed and the interquartile range (IQR) was used to remove outliers. ECHO, Welch’s two-sample T-tests, and Hedges’ g were used to determine the statistical significance of the differences between the average cellular pixel intensities^42^. We ran a Hedges’ g since it is an appropriate statistical analysis to measure effect with samples of varying sample size. A Hedges’ g value above 0.2 shows a small effect, above 0.5 shows a medium effect, and above 0.8 shows a large effect; therefore, a large Hedges’ g confirms significance whereas a small Hedges’ g negates significance. *All raw phagocytosis data of cellular pixel intensities will be uploaded to Mendeley Data upon publication*.

### ECHO Analysis

ECHO version 4.1 was used to analyze all transcriptome, proteome, PG concentrations, and phagocytosis data that had samples gathered over circadian time^42^. All data were free run with the smooth data and linear detrend data options selected. Transcriptome data is reported in normalized transcripts per million and the proteome data is reported in normalized spectral counts. Transcriptome and Proteome data analyzed in ECHO has an x-axis of 0 to 46 corresponding to the number of samples taken in chronological order, which when added to 16 equals the post shock time of each sample. In order to calculate zeniths for the transcriptome and proteome data, the hours shifted from the ECHO analysis was added to 16 to get the post shock time, which was then converted to CT time using Keller et al., 2009^41^. PG level analysis included all four replicates per time point. For the phagocytosis data, the average of the replicates was analyzed at each time point.

### Liquid Chromatography-Tandem Mass Spectrometry

Using synchronized macrophages (as described above) starting at PS16 every 4 h for 24 h (PS16, PS20, PS24, PS28, PS32, PS36, and PS40), macrophages, in quadruplicate, were washed once with PBS, and removed from the plate using Cell Stripper (Corning 25-056-CI). Macrophages were centrifuged at 400 *g* for three minutes, the supernatant was aspirated, the cells were resuspended in 250 µL PBS, and flash frozen. Additionally, at PS16, PS20, PS28, and PS36, spent media from each culture dish was centrifuged at 400 x g for three minutes to remove any particulates and flash frozen. Finally, after macrophage removal, at PS16, PS20, PS28, and PS36, the culture dish was scraped using a sterile rubber policeman, the dish was flushed with 1 ml of PBS, and the resultant sample was flash frozen to sample the ECM.

To analyze total cellular levels of PGs using LC-MS/MS, macrophage pellets were treated with 100 μl BugBuster 10× protein extraction reagent (MilliporeSigma) and transferred into 1 mL microcentrifuge tubes, which were then sonicated at room temperature for 20 min. After that, 300 μl digestion buffer (50 mM ammonium acetate, 2 mM calcium chloride) was added. To analyze total supernatant and ECM levels of PGs using LC-MS/MS, 400 μl of clarified supernatant/ECM scrapings were loaded onto a 3 KDa spin column (Millipore Sigma UFC500396), washed with distilled water, and then the upper solution was aspirated and mixed with 300 μl digestion buffer. In all cases, recombinant heparin lyase I, II, III and chondroitin lyase ABC (10 mU each) (prepared by the Linhardt lab) were added to the 300 μl of digestion buffer and incubated at 37°C overnight. Digestion reactions were terminated by passing the sample through a 3 KDa MWCO spin column to eliminate the lyases. The 3 KDa MWCO spin columns were washed twice with 300 μl distilled water and the filtrate was lyophilized. Lyophilized samples were AMAC-labeled by incubating them at room temperature for 10 min with 10 μl of 0.1 M AMAC in DMSO/acetic acid (17/3,V/V), after which 10 μl of 1 M aqueous NaBH_3_CN was added and the mixture was incubated for 1 h at 45°C. The resultant samples were centrifuged at 13,200 rpm for 10 min. Finally, each supernatant was collected and stored in a light resistant container at room temperature until LC-MS/MS analysis.

LC was performed on an Agilent 1200 LC system at 45°C using an Agilent Poroshell 120 ECC18 (2.7 μm, 3.0 × 50 mm) column. Mobile phase A (MPA) was a 50 mM ammonium acetate aqueous solution, and mobile phase B (MPB) was methanol. The mobile phase passed through the column at a flow rate of 300 μl/min. The gradient was 0-10 min, 5-45% B; 10-10.2 min, 45-100%B; 10.2-14 min, 100%B; 14-22 min, 100-5%B. A triple quadrupole mass spectrometry system equipped with an ESI source (Thermo Fisher Scientific, San Jose, CA) was used as a detector. The online MS analysis was in the multiple reaction monitoring (MRM) mode. MS parameters: negative ionization mode with a spray voltage of 3,000 V, a vaporizer temperature of 300°C, and a capillary temperature of 270°C. The conditions and collision energies for all of the disaccharides MRM transitions are listed in Table S6. Sample measurements for the cell pellet were normalized in nanograms of PGs per cell (ng/cell) using a CyQUANT Assay to quantify the number of cells used per pellet as described in the methods. Sample measurements from the ECM and spent media samples were calculated in nanograms per milliliter (ng/ml). *All HSPG and CSPG raw data generated from this experiment will be uploaded to Mendeley Data upon publication*.

### CyQUANT Assay

Due to the varying amount of protein present in a cell over time, we normalized the HSPG and CSPG cell pellet findings to the number of cells present in each sample. The cells were quantified using a CyQUANT Cell Proliferation assay (Invitrogen C7026). This kit was used to lyse 100 μl of the cells stripped from each plate and stain the nucleic acids with a fluorescently labeled dye (included in kit cited above), which was then read by a plate reader at an excitation wavelength of 480 nm and emission wavelength of 520 nm. The concentration of cells was then back calculated using a standard curve made by measuring known amounts of the primary murine macrophages. This standard curve was constructed per the manufacturer’s instructions (Invitrogen C7026 manual). The cells were diluted to a known amount using a BioRad TC20 automated cell counter, lysed, dyed, and read on a Tecan Infinite M1000 Pro plate reader at an excitation wavelength of 480 nm and emission wavelength at 520 nm. The number of cells from each sample was first calculated using the equation of the standard curve (R^2^=0.9938) and then multiplied by the dilution factor to calculate the total number of cells in each cell pellet.

### Production of Heparinases

Heparinases were prepared as has been previously reported^63–65^. All heparinases used were purified away from endotoxins using an Endotoxin Removal Kit from GenScript (L00338)^66^.

