## Supplemental Figures for "Circadian Control of Heparan Sulfate Levels Times Phagocytosis of Amyloid Beta Aggregates"

### Supplemental Information

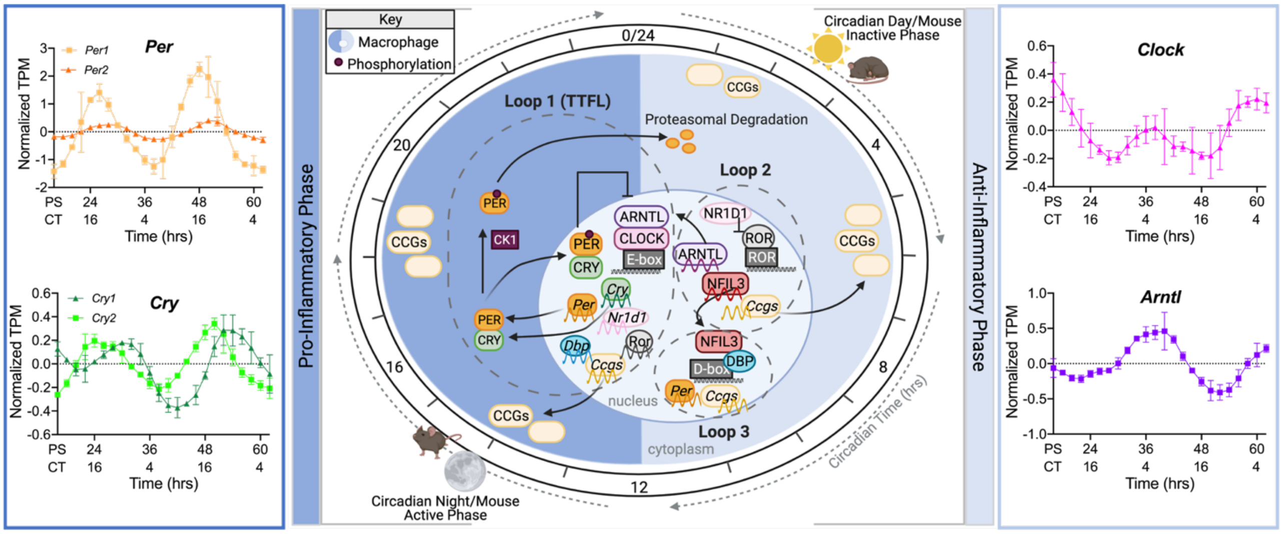


<u>Supplemental Figure 1: The murine circadian clock times macrophage physiology.</u> The macrophage circadian rhythm is imposed by a conserved, core molecular clock that imparts control of key gene and protein expression through a transcription/translation negative feedback loop (TTFL). In murine macrophages, pictured as the blue cell above, the positive arm of the clock, in the core circadian timekeeper (Loop 1), is made up of ARNTL and CLOCK, which binds to the E-box promoter to turn on the creation of the negative arm protein (PER, CRY), auxiliary loop proteins (NR1D1, DBP, ROR), and *clock-controlled genes* (*ccgs*). The negative arm proteins then dimerize in the cytoplasm and are temporally phosphorylated by CK1 (also known as Csnk1a1 in mice), until they migrate back into the nucleus and inhibit their own transcription by inhibiting ARNTL and CLOCK, the negative arm is then ubiquinated and degraded, restarting the cycle. Two auxiliary loops to the circadian clock, Loop 2 and Loop 3, impart further fine-tuning of circadian regulation. In Loop 2, the cycle starts when ROR binds to the ROR promoter sequences to turn on expression of *arntl*, *nfil3*, and other *ccgs*. This loop is inhibited when the positive arm activates transcription of NR1D1, which inhibits transcription in Loop 2. Loop 3 is activated via the DBP binding to the D-box promoter and turns on expression of PER and CCGs. This expression is in turn inhibited by the expression of NFIL3 from Loop 2. The surrounding gene expression graphs show the timing of expression of key clock proteins in Post Shock (PS) time and Circadian Time (CT) using transcriptome data from Collins et al., 2020 [S1]. The murine clock pathway was adapted from Curtis et al., 2014 [S2].

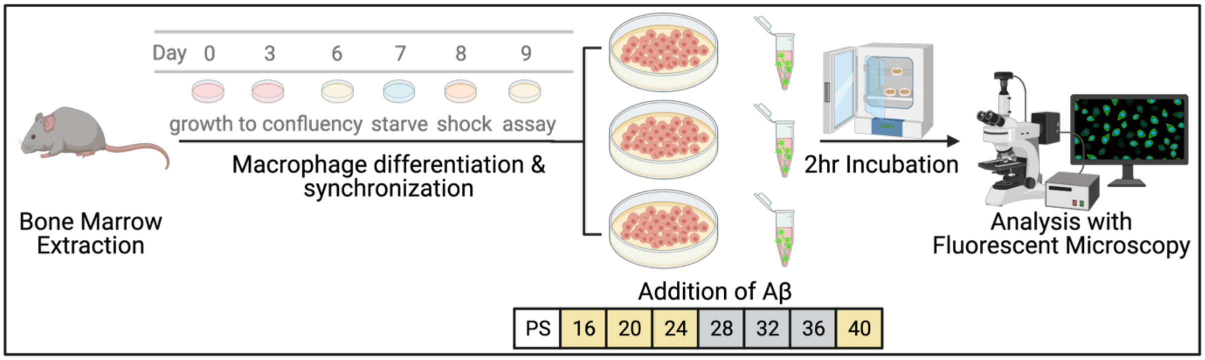


Supplemental Figure 2: Schematic of the macrophage phagocytosis assay protocol. Our procedure began with the extraction of bone marrow monocytes from PER2::LUC mice. Extracted bone marrow monocytes were treated with mCSF with fresh media (10% FBS) added every 3 days and these bone marrow derived macrophages were grown to confluency. After seven days, macrophages were transferred to media without serum (0%FBS) for 24 hours before being transferred to serum shock media (50% FBS) for 2 hours, in order to synchronize their clocks. Synchronized macrophages were transferred into fresh media (10% FBS) and, starting at 16 hours post shock (PS 16), samples were treated in triplicate with Aβ (for 2 hours) every four hours for 24 hours. Macrophages were then harvested, fixed using formalin, and imaged using fluorescence microscopy. PS hours as represented in yellow for the active and grey for the inactive phase of the mice as reported previously [S1].

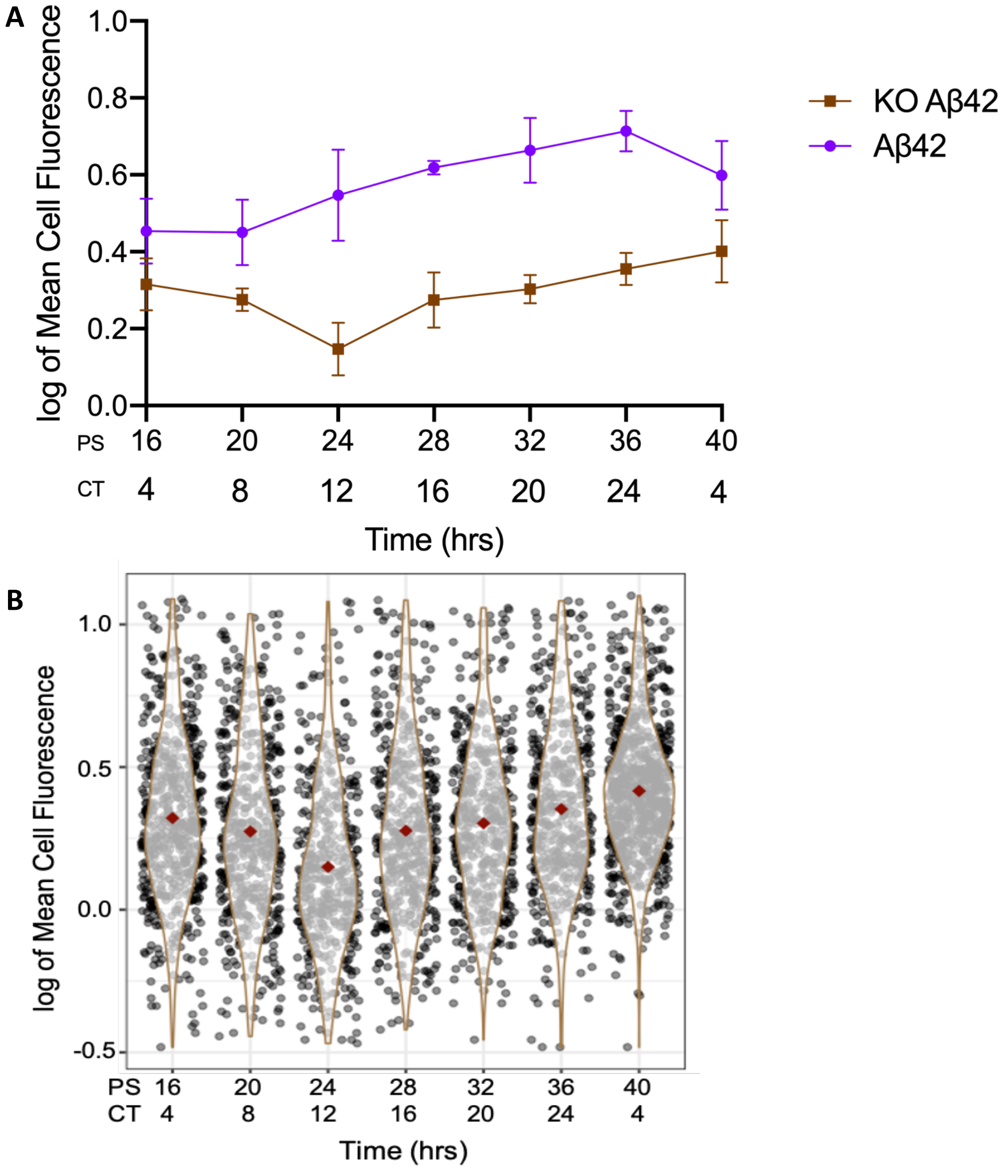


<u>Supplemental Figure 3: Aβ42 Phagocytosis Oscillates with a Circadian Period.</u> A) The log of mean cell fluorescence of Aβ42 phagocytosis in PER2::LUC cells (purple circles), and in PER1$^{-/-}$PER2$^{-/-}$ knockout cells (brown squares) plotted over time. Error bars represent the standard deviation. B) Violin plot of Aβ42 phagocytosis in PER1$^{-/-}$PER2$^{-/-}$ knockout cells represented in logged mean cell fluorescence values plotted against post shock (PS) and circadian time (CT). Grey dots represent a single cell measurement, brown violin plot lines represent the range of the data, and red diamonds indicate the mean measurement of each time point. CT is in reference to the comparison of PER2 expression from Keller et al., 2009 [S3] compared to PER2 expression from our cells. All data shown was performed in triplicate.

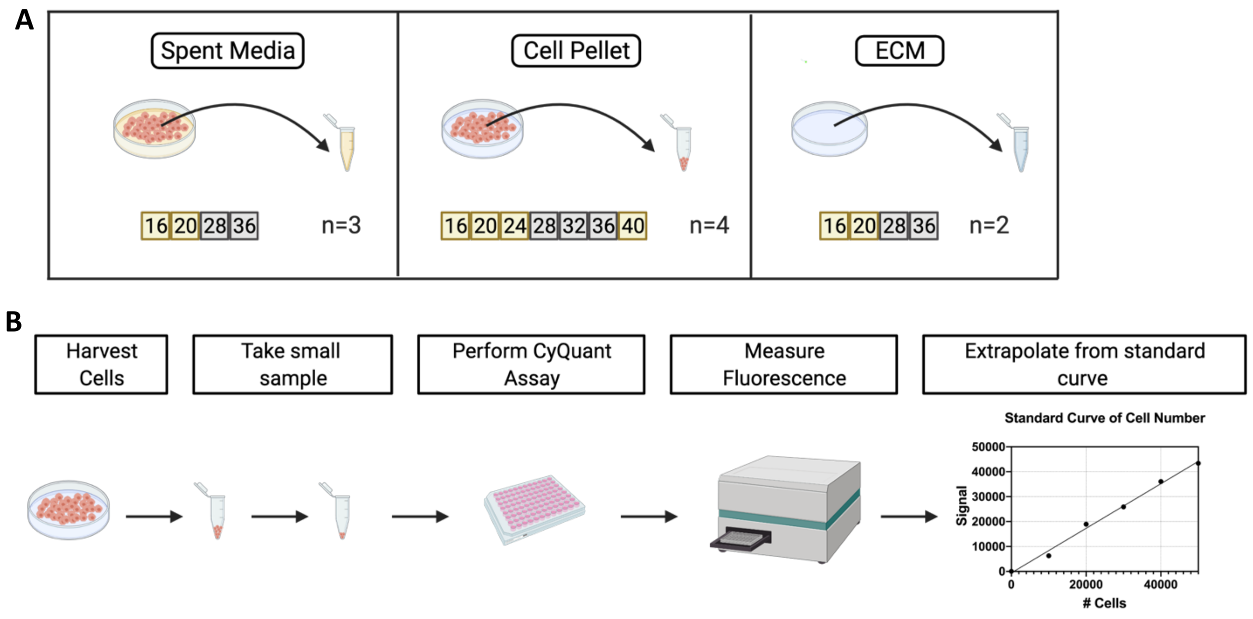


<u>Supplemental Figure 4: Schematic of the quantification of PGs in BMDMs.</u> (A) BMDMs were collected at PS 16, 20, 24, 28, 32, and 36, in quadruplicate. Spent media from these cell pellets was collected at PS 16, 20, 28, and 36, in triplicate. Concurrently, ECM samples were collected in duplicate at post shock times 16, 20, 28, and 36. (B) The above samples were analyzed by LC-MS/MS and normalized using data from a CyQuant assay that calculated total cellular DNA by UV-VIS spectroscopy and an interpolated standard curve (standard curve in inset from actual macrophage data).

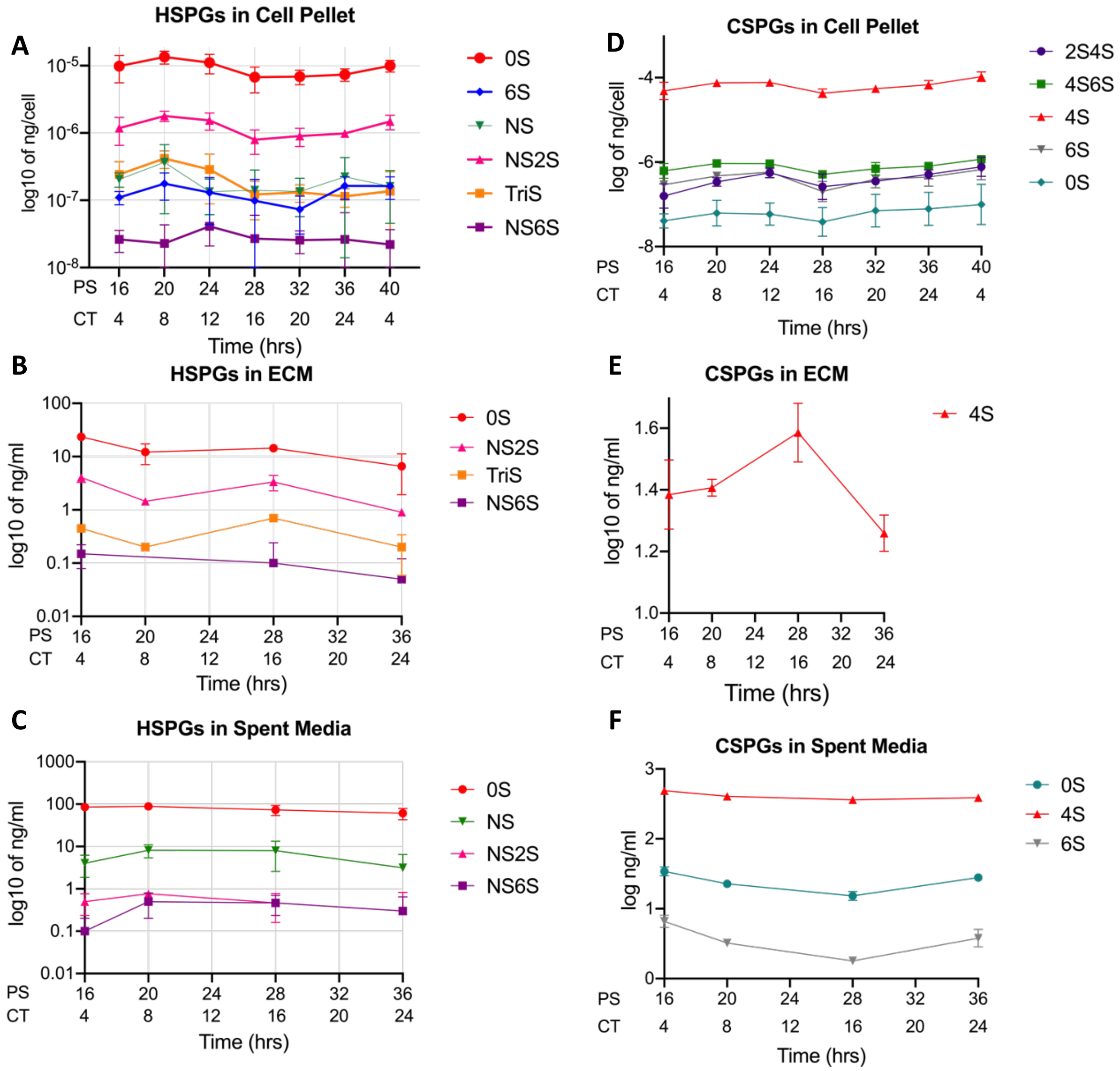


<u>Supplemental Figure 5: HSPG and CSPG levels oscillate over circadian time in murine macrophages.</u> (A) The level of each HSPG sulfation type in macrophage cells (n=4) plotted against PS and CT time. (B) The level of each HSPG sulfation type in macrophage ECM scrapings (n=2) plotted against PS and CT time. (C) The level of each HSPG sulfation type in macrophage spent media cultures (n=3) plotted against PS and CT time. (D) The level of each CSPG sulfation type in macrophage cells (n=4) plotted against PS and CT time. (E) The level of each CSPG sulfation type in macrophage ECM scrapings (n=2) plotted against PS and CT time. (F) The level of each CSPG sulfation type in macrophage spent media cultures (n=3) plotted against PS and CT time. n = number of replicates, error bars are represented by the standard deviation of the replicates.

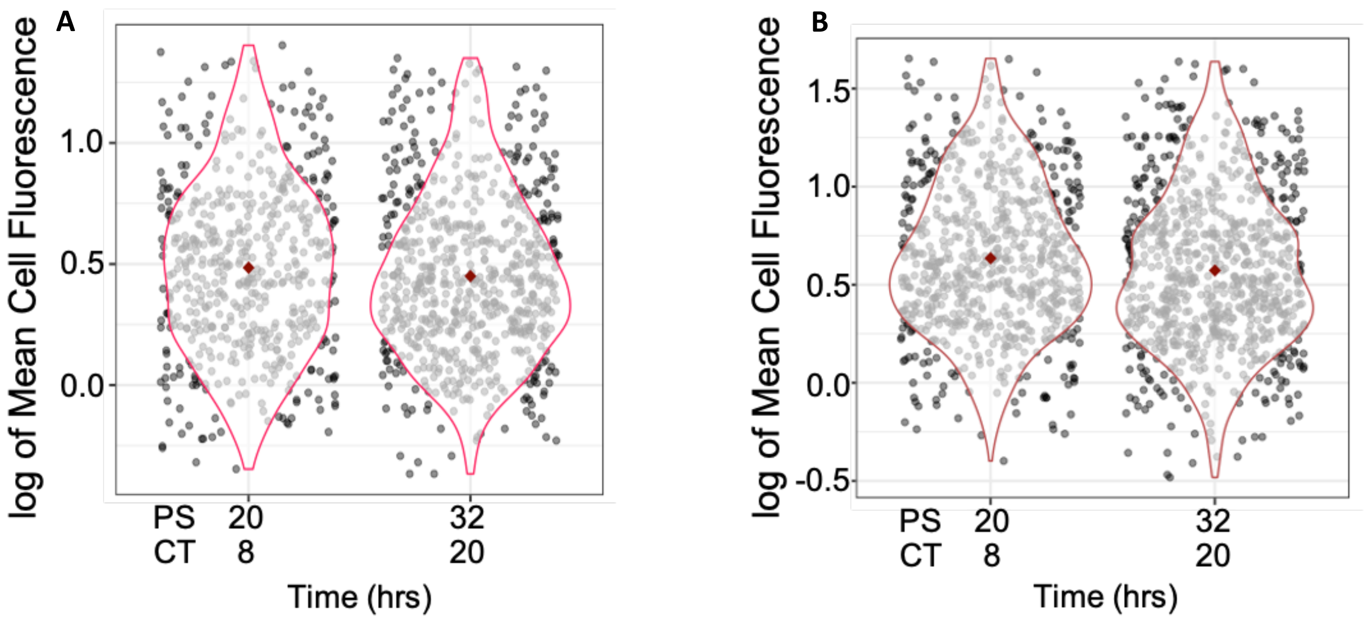


Supplemental Figure 6: Heparinases I and III differentially disrupt the phagocytosis of Aβ42. Violin plot of Aβ42 phagocytosis in the present of A) Heparinase I and B) Heparinase III represented in logged mean cell fluorescence values plotted against PS and CT. Grey dots represent a single cell measurement, A) red and B) maroon violin plot lines represent the range of the data, and red diamonds indicate the mean measurement of each time point. CT is in reference to the comparison of PER2 expression from Keller et al., 2009 [S3] compare to PER2 expression from our cells.

Supplemental Table 1: ECHO analysis of the Aβ42 phagocytosis assays.

Supplemental Table 2: ECHO analysis of the transcripts and proteins in the PG pathway.

Supplemental Table 3: ECHO analysis of lipid receptors and cell cycle proteins.

Supplemental Table 4: ECHO analysis of the HSPG and CSPG levels in the macrophage cell pellet.

Supplemental Table 5: Heparinase Concentrations and Activities.

Supplemental Table 6: Conditions and collision energies for the disaccharide MRM transitions.
